# Re-expression of SynGAP Protein in Adulthood Improves Translatable Measures of Brain Function and Behavior in a Model of Neurodevelopmental Disorders

**DOI:** 10.1101/474965

**Authors:** Thomas K. Creson, Camilo Rojas, Ernie Hwaun, Thomas Vaissiere, Murat Kilinc, J. Lloyd Holder, Jianrong Tang, Laura Lee Colgin, Courtney A. Miller, Gavin Rumbaugh

## Abstract

**Background:** Neurodevelopmental disorder (NDD) risk genes have pleiotropic biological functions, such as control over both developmental and non-developmental processes that influence disease-related phenotypes. Currently, it remains unclear how developmental versus non-developmental processes influence the duration and/or effectiveness of permissive treatment windows for NDDs. *SYNGAP1* haploinsufficiency causes an NDD defined by autistic traits, cognitive impairment, and epilepsy. *Syngap1* heterozygosity in mice disrupts a developmental critical period, and, consistent with this, certain behavioral abnormalities are resistant to gene therapy initiated in adulthood. However, the *Syngap1* endophenotype is extensive and this protein has diverse cell biological functions. Therefore, SynGAP pleiotropy may influence the permissive treatment window for previously untested disease-relevant phenotypes.

**Methods:** A whole-body gene restoration technique was used to determine how restoration of SynGAP protein in adult heterozygous mice impacted previously untested phenotypes, such as memory, seizure susceptibility, systems-level cortical hyperexcitability, and hippocampal oscillations linked to mnemonic processes.

**Results:** Adult restoration of SynGAP protein in haploinsufficient mice reversed long-term contextual memory deficits and behavioral measures of seizure susceptibility. Moreover, SynGAP re-expression in adult mice eliminated brain state-dependent, patient-linked paroxysmal interictal spiking and increased the amplitude of hippocampal theta oscillations.

**Conclusions:** SynGAP protein in the mature brain dynamically regulates neural circuit function and influences disease-relevant phenotypes. The impact of these findings is that treatments targeting certain debilitating aspects of *SYNGAP1-*related disorders may be effective throughout life. Moreover, the efficacy of experimental treatments for *SYNGAP1* patients may be quantifiable through changes in species-conserved, state-dependent pathological electroencephalogram signals.

## Introduction

Neurodevelopmental disorders (NDDs), including intellectual disability (ID) and autism spectrum disorder (ASD), are common and often result in treatment-resistant cognitive and behavioral impairments[1]. Historically, the pathoneurobiology underlying NDD symptomology was thought to arise from impaired brain development. However, studies in animal models of genetic risk factors causally-linked to syndromic NDDs have shown improvement in meaningful measures of brain function and behavior in response to adult-initiated therapeutic interventions[2-4]. These findings suggest that not all phenotypic consequences of NDDs arise through impaired neurodevelopment. This has led to the idea that therapeutic intervention may be beneficial in adult NDD patients with fully mature brains[5, 6]. The potential impact of adult initiated treatments cannot be overstated, as there are no currently effective treatments for the most debilitating aspects of these disorders. Thus, as new treatments are developed, hope remains that adults with NDDs could benefit from these emerging therapeutic strategies.

Over the past decade, hundreds of new genes have been linked to NDDs[7]. Large-scale exome sequencing projects in children with classically undefined and sporadic NDDs have identified a pool of high-risk genes that are now causally-linked to these disorders[8, 9]. Some of these newer NDD genes account for a significant fraction of total cases. Given that the list of high-impact genes has considerably expanded over the last decade, it is unknown to what extent the approach of adult reversibility applies to disease-relevant phenotypes caused by pathogenicity of these newly discovered NDD risk factors.

*SYNGAP1* is a recently discovered high-risk NDD gene[10-12]. It is causally linked to a range of sporadic NDDs, including ID[8, 9, 12, 13], ASD[14, 15], severe epilepsy[16, 17] and schizophrenia[18]. *De novo* nonsense variants in *SYNGAP1* resulting in haploinsufficiency lead to a relatively common genetically-defined form of ID with epilepsy (MRD5; OMIM#603384) that may explain up to 1% of ID cases[19, 20]. MRD5 patients express moderate-to-severe intellectual disability (IQ <50), have severely delayed language development, and express some form of epilepsy and/or abnormal brain activity, with these manifestations appearing first in early childhood[19-21]. Most MRD5 patients have severe loss-of-function *SYNGAP1* variants in one allele[21] that leads to reduced protein expression or function (i.e. genetic haploinsufficiency).

*Syngap1* heterozygous knockout mice (Hets), which have ~50% reduction in SynGAP protein levels[22], offer both construct and face validity for MRD5[23]. *Sygnap1* heterozygosity disrupts a developmental critical period that is essential for the assembly and excitatory balance of developing forebrain circuits[22, 24-26]. Prior studies have shown that reduced *Syngap1* expression during this developmental critical period prevents the normal emergence of certain behavioral responses and cognitive functions because altered measures of working memory, locomotor activity, and anxiety were sensitive to neonatal reversal of *Syngap1* pathogenicity[24], but resistant to similar approaches performed in adulthood[22]. However, knocking out SynGAP protein in the adult hippocampus causes memory impairments[27], suggesting that *Syngap1* also has unique, non-developmental functions in adult neurons that support cognitive functions. Importantly, *Syngap1* mice have an extensive endophenotype[22, 26, 28, 29], though only a small subset of individual phenotypes observed in this line have been tested for sensitivity to adult reversal of genetic pathogenicity[22]. Based on these past results, we determined if previously untested elements of the broader *Syngap1* behavioral endophenotype are sensitive to a method of adult gene restoration that restores pathologically low levels of SynGAP protein. Moreover, we also searched for neurophysiological correlates of behavioral alterations in *Syngap1* animals to gain insight into how protein re-expression in mature animals may improve brain function.

## Results

To date, only spontaneous alternation has been used to determine the impact of adult reversal of *Syngap1* pathogenicity on changes to cognitive function[22]. This measure of memory was not improved after adult reversal *Syngap1* Hets. We were, therefore, interested in identifying a distinct behavioral paradigm reflective of impaired Het cognitive function so that it could be tested for sensitivity to adult reversal of pathologically low SynGAP protein levels. We found that two different *Syngap1* Het lines exhibit a remote memory impairment when tested 26-30 days after contextual fear training **(Fig. 1a, b, e, f).** Interestingly, *Syngap1* Hets have normal contextual memory when tested just one day after training **(Fig. 1c)**[28, 29], indicating that either systems memory consolidation or remote memory retrieval is disrupted by germline expression of *Syngap1* pathogenicity. Surprisingly, we found that retrieval 1d after training prevented 26-30d remote memory deficits in *Syngap1* Hets **(Fig. 1c)**.

**Figure 1:**
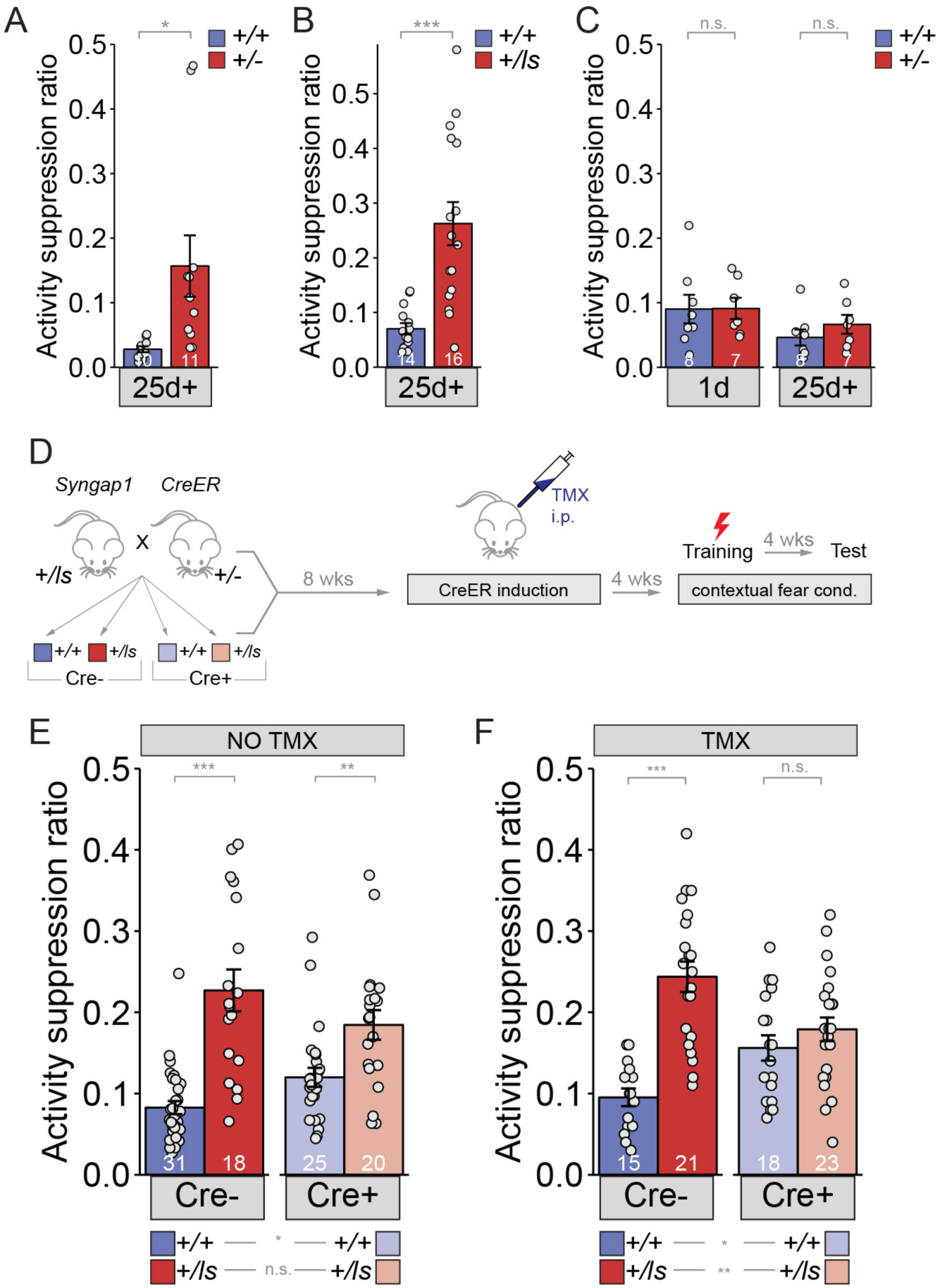
Long-term memory can be improved in adult mice with *Syngap1* pathogenicity. (A) *Syngap1*^*+/+*^ and *Syngap1*^*+/-*^ mice were trained in the remote contextual fear conditioning paradigm and tested one month later for activity suppression levels. Activity of the *Syngap1*^*+/-*^ group was suppressed significantly less than that of the *Syngap1*^*+/+*^ group indicating compromised remote memory for the mutant group. Unpaired *t* test (t(19)=-2.567, p=.019). **(B)** *Syngap1*^*+/+*^ and *Syngap1*^*+/Is*^ mice were trained in the contextual fear conditioning paradigm and tested one month later for activity suppression levels. Activity of the *Syngap1*^*+/Is*^ group was suppressed significantly less than that of the *Syngap1*^*+/+*^ group indicating compromised remote memory for the mutant group. Wilcoxon rank sum test W=19, p=2.82E-5 **(C)** *Syngap1*^*+/+*^ and *Syngap1*^*+/-*^ mice were tested, firstly, 1d after training, followed by another testing one month later. Activity suppression levels were not significantly different between the groups for either testing (unpaired t test,1 day t(13)=- 0.033, p=0.974; 26 days t(13)=-1.068, p=0.305). (**D)** Experimental schematic depicting the breeding strategy for generation of ere-inducible *Syngap1*^*Cre+;+/ls*^ mice and Cre induction with tamoxifen treatment for restoration of *Syngap1* expression and subsequent remote fear conditioning testing. **(E-F)** *Syngap1*^*Cre-;+/+*^, *Syngap1*^*Cre-;+/ls*^, *Syngap1*^*Cre+;+/+*^, and *Syngap1*^*Cre+;+/ls*^ mice were run in the remote contextual fear conditioning paradigm without (E) and with (F) TMX administration. Activity suppression values from mice without TMX administration (**No TMX**) were assessed (2-factor ANOVA: [add F values throughout just for main effects] Main Effects-Cre F(1,90)=.030, p=.864, Genotype F(1,91)=46.78, p=9.28E-10, Interaction F(1,91)=6.81, p=.011. Pairwise comparisons-*Syngap1*^*Cre-;+/+*^ vs *Syngap1*^*Cre-;+/ls*^ (p=1.58E-9), *Syngap1*^*Cre+;+/+*^ vs *Syngap1*^*Cre+;+/ls*^ (p=.004), *Syngap1*^*Cre-;+/+*^ vs *Syngap1*^*Cre+;+/+*^ (p=.059), *Syngap1^Cre-;+/ls^ vs Syngap1^Cre+;+/ls^* (p=.074); **With TMX administration** (2-factor ANOVA: Main Effects-Cre F=(1,73)=.019, p=.891, Genotype F(1,73)=27.49, p=1.48E-6, Interaction F(1,73)=14.75, p=2.59E-4. Pairwise comparisons- *Syngap1*^*Cre-;+/+*^ vs *Syngap1*^*Cre-;+/ls*^ (p=2.98E-8), *Syngap1*^*Cre+;+/+*^ vs *Syngap1*^*Cre+;+/ls*^ (p=.307), *Syngap1^Cre-;+/+^ vs Syngap1^Cre+;+/+^* (p=.017), *Syngap1^Cre-;+/ls^ vs Syngap1^Cre+;+/ls^* (p=.003).

Retrieval induced strengthening suggested that impairments underlying the remote memory deficit are not hard-wired in Hets and may, therefore, be sensitive to adult re-expression of SynGAP protein. To test this, we crossed *Syngap1* Lox-stop Het mice to hemizygous mice expressing a tamoxifen (TMX)-inducible form of Cre recombinase[22]. We then trained adult offspring resulting from this cross in contextual fear conditioning and tested memory ~1 month after treatments with or without TMX **(Fig. 1d)**. In TMX(-) mice, both Cre (-) and Cre(+) *Syngap1* Hets were significantly different compared to corresponding *Syngap1* WT controls. Importantly, there was no difference between Cre(-) and Cre(+) *Syngap1* Hets **(Fig. 1e)**. However, TMX-treated Cre(+) *Syngap1* Het mice (*i.e. mice with Syngap1 pathogenicity reversed starting at PND60; *Supplementary Figure 1*)[22]* were no different from corresponding WT *Syngap1* mice **(Fig. 1f)**. Moreover, the activity suppression ratio in these mice was significantly reduced compared to Cre(-) *Syngap1* Hets (i.e. mice with preserved *Syngap1* pathogenicity), consistent with adult reversal of the remote memory deficit.

MRD5 patients *and Syngap1* Hets both express electrographic and behavioral seizures [21, 26, 30]. Seizures in many MRD5 patients are medically refractory. To determine the permissive treatment window for seizure-related behaviors in *Syngap1* mice, we explored how SynGAP re-expression impacted measures of seizure susceptibility in adult *Syngap1* Hets [22, 26]. In TMX(-) mice, both Cre (-) and Cre(+) *Syngap1* Lox-Stop Hets were significantly different compared to corresponding *Syngap1* WT mice in the first two stages of seizure **(Fig. 2a)**, which replicates our previous findings in adult constitutive *Syngap1* Heterozygous knockout mice[26]. Importantly, there was no difference between Cre(-) and Cre(+) *Syngap1* Lox-Stop Hets, indicating that Cre remained inactive in Cre(+)TMX(-) mice. However, similar to what we observed in fear conditioning studies, TMX-treated Cre(+) *Syngap1* Het mice were no different from Cre(+) WT *Syngap1* mice **(Fig. 2b)**. Furthermore, the time to seizure in Cre(+) *Syngap1* Het mice was significantly increased compared to Cre(-) *Syngap1* Hets in the two stages of the test that show impairments. These data demonstrate that impaired excitability linked to seizure susceptibility caused by *Syngap1* pathogenicity can be improved in adulthood after SynGAP re-expression.

**Figure 2:**
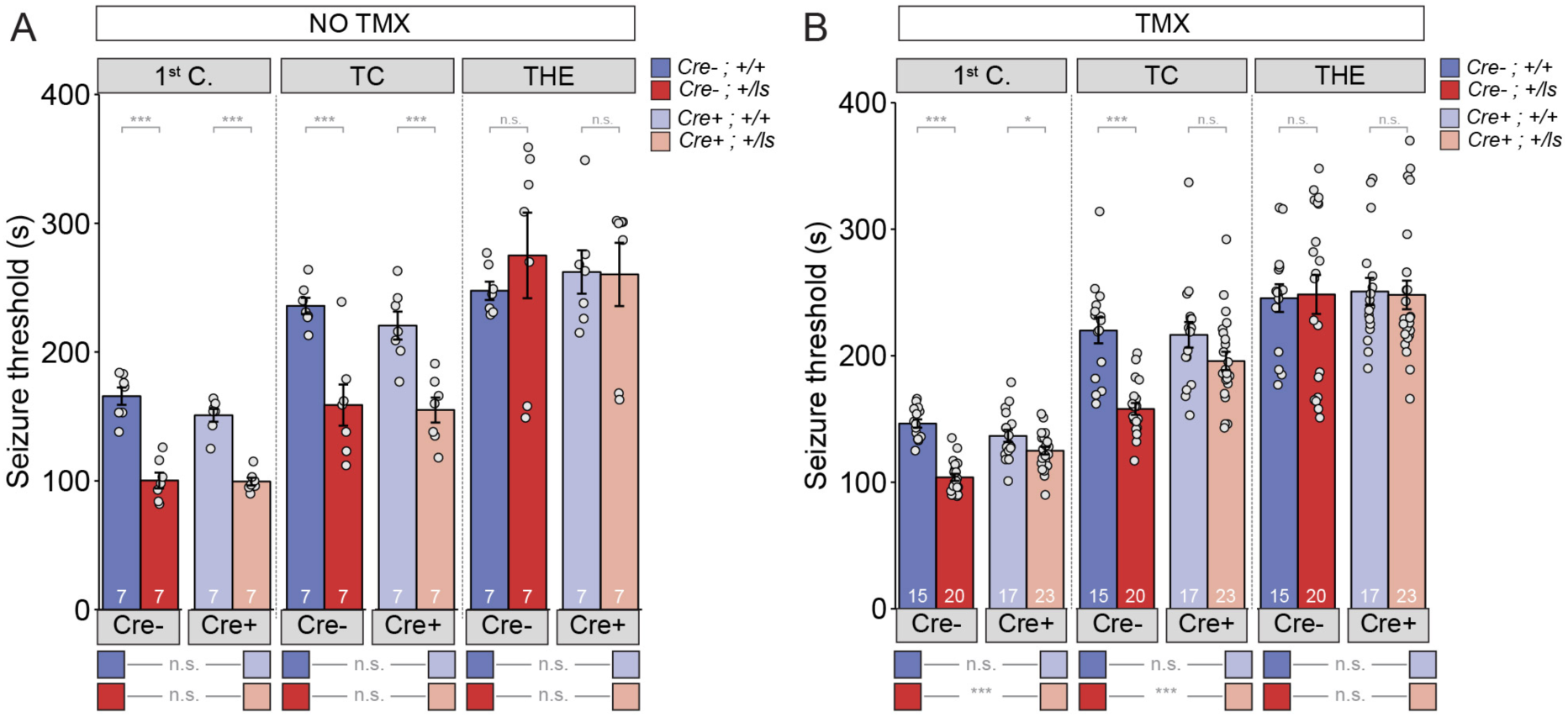
Seizure threshold, but not sensory-motor gating, is improved after adult restoration of SynGAP expression. (A) *Syngap^Cre-;+/Is^* and *Syngap^Cre+;+/Is^* mice exhibit hyperexcitability in two of the three events without Cre activation (**No TMX**) Main effects-1^st^ clonus: Cre F(1,24)=2.13, p=.157, Genotype F=117.73, p=9.75E-11, Interaction F(1,24)=1.69, p=.206); TC: Cre F(1,24)=722, p=.404, Genotype F(1,24)=40.05, p=1.53E-6), Interaction F(1,24)=.257, p=.617); THE: Cre F(1,24)=9.99E-6, p=.998, Genotype F(1,24)=.320, p=.577), Interaction F(1,24)=.420, p=.523). Pairwise comparisons-1^st^ clonus: *Syngap1*^*Cre-;+/+*^ vs *Syngap1*^*Cre-;+/Is*^ (p=8.72E-9)), *Syngap1*^*Cre+;+/+*^ vs *Syngap1*^*Cre+;+/Is*^ (p=5.51E-7), *Syngap1^Cre-;+/+^ vs Syngap1^Cre+;+/+^*(p=.063), *Syngap1*^*Cre-;+/Is*^ vs *Syngap1*^*Cre+;+/Is*^ (p=.911). T/C: *Syngap1*^*Cre-;+/+*^ vs *Syngap1*^*Cre-;+/Is*^p=6.34E-5), *Syngap1*^*Cre+;+/+*^ vs *Syngap1*^*Cre+;+/Is*^ (p=3.93E-4), *Syngap1^Cre-;+/+^ vs Syngap1^Cre+;+/+^* (p=.347), *Syngap1^Cre-;+/Is^ vs Syngap1^Cre+;+/Is^*(p=.811). THE: *Syngap1*^*Cre-;+/+*^ vs *Syngap1*^*Cre-;+/Is*^(p=.399), *Syngap1^Cre+;+/+^ vs Syngap1^Cre+;+/Is^* (p=.954), *Syngap1^Cre-;+/+^ vs Syngap1^Cre+;+/+^* (p=.652), *Syngap1^Cre-;+/Is^ vs Syngap1^Cre+;+/Is^*(p=.649). **(B)** *Syngap^Cre+;+/Is^* mice exhibit thresholds comparable to those of *Syngap^Cre-;+/Is^* mice after Cre activation (TMX) in two of the three events Main effects-1^st^ clonus: Cre F(1,71)=2.59, p= .112; Genotype F(1,71)=58.328, p= 7.86E-11, Interaction F=1(1,71)=18.84 p=4.62E-5; TC: Cre F(1,71)=4.53, p=.037, Genotype F(1,71)=26.15, p=2.57E-6, Interaction F(1,71)=6.50, p=.013; THE: Cre F(1,71)=.037, p=.847, Genotype F(1,71)=1.15E-5, p=.997, Interaction F(1,71)=.049, p=.826. Pairwise comparisons-1^st^ clonus: *Syngap1*^*Cre-;+/+*^ vs *Syngap1*^*Cre-;+/Is*^(p=6.96E-12), *Syngap1*^*Cre+;+/+*^ vs *Syngap1*^*Cre+;+/Is*^ (p=.018), *Syngap1^Cre-;+/+^ vs Syngap1^Cre+;+/+^*(p=.076), *Syngap1^Cre-;+/Is^ vs Syngap1^Cre+;+/Is^*(p=2.13E-5). T/C: *Syngap1*^*Cre-;+/+*^ vs *Syngap1*^*Cre-;+/Is*^(p=1.52E-6), *Syngap1*^*Cre+;+/+*^ vs *Syngap1*^*Cre+;+/Is*^(p=.065), *Syngap1^Cre-;+/+^ vs Syngap1^Cre+;+/+^* (p=.782), *Syngap1^Cre-;+/Is^ vs Syngap1^Cre+;+/Is^* (p=001). THE: *Syngap1*^*Cre-;+/+*^ vs *Syngap1*^*Cre-*^(p=.878), *Syngap1*^*Cre+;+/+*^ vs *Syngap1*^*Cre+;+/Is*^ (p=.874), *Syngap1^Cre−^;+/+ vs Syngap1^Cre+;+/+^* (p=.785), *Syngap1^Cre-;+/Is^ vs Syngap1^Cre+;+/Is^*(p=. 983).

Given the behavioral improvements in response to SynGAP re-expression in adult Cre(+)/Lox-Stop Het mice, we explored how this approach impacts neurophysiological correlates of seizure and memory in these mice. Prior studies have shown through EEG recordings that conventional *Syngap1* Heterozygous knockout mice display high-amplitude interictal spiking events [26]. Interictal spikes (IIS) are pathological electrical events that reflect seizure susceptibility in patients and may share common mechanisms with ictal events [31]. We reasoned that because behavioral seizure susceptibility was reversed after SynGAP re-expression in Cre(+)/Lox-Stop Hets, IIS events may also be ameliorated by this therapeutic approach. However, neurophysiological studies have not been performed in *Syngap1* Lox-Stop mice. Therefore, it is unknown if IIS events are present in this line. We performed *in vivo* neurophysiological recordings in both Cre(+) WT and Cre(+) Lox-Stop Het mice. In addition to left and right sub-cranial EEG electrodes, a depth electrode was lowered into CA1 to acquire a hippocampal local field potential (*h*LFP). We carried out chronic recordings in these animals and the study consisted of two recording phases. Phase 1 was geared toward identifying putative genotype-dependent differences in neurophysiological signals between Cre(+) WT and Cre(+) Lox-Stop mice. Phase II, in which all animals in Phase I underwent a procedure to induce Cre recombinase immediately following Phase I studies, was geared toward determining how putative neurophysiological abnormalities in each animal were impacted by restoration of SynGAP levels in Cre(+) Lox-Stop Het mice. In Phase I recordings, we observed frequent high amplitude IIS events that generalized across the three recording channels in Lox-Stop mice **(Fig. 3A; Supplemental Figure 2).** This finding demonstrates that IIS is a reproducible phenotype in mice with *Syngap1* haploinsufficiency. We quantified the frequency of IIS events during wakefulness in each genotype and found that spiking events were ~50 fold more frequent in mutants compared to WT controls **(Fig. 3B)**. We also observed that high-amplitude IIS events in Lox-Stop animals exhibited state-dependence during Phase I recordings, with an ~18-fold higher frequency of these events observed during sleep compared to wakefulness **(Fig. 3A, C)**. We took advantage of the ability to re-express SynGAP protein in Lox-Stop mice to investigate how this strategy impacted IIS events. Remarkably, recordings during Phase II revealed that SynGAP re-expression in adult Lox-Stop mice eliminated IIS activity during wakefulness and sleep **(Fig. 3A-B).** These data demonstrate that SynGAP protein dynamically regulates cellular mechanisms that govern systems-level excitability in the mature mouse forebrain.

**Figure 3:**
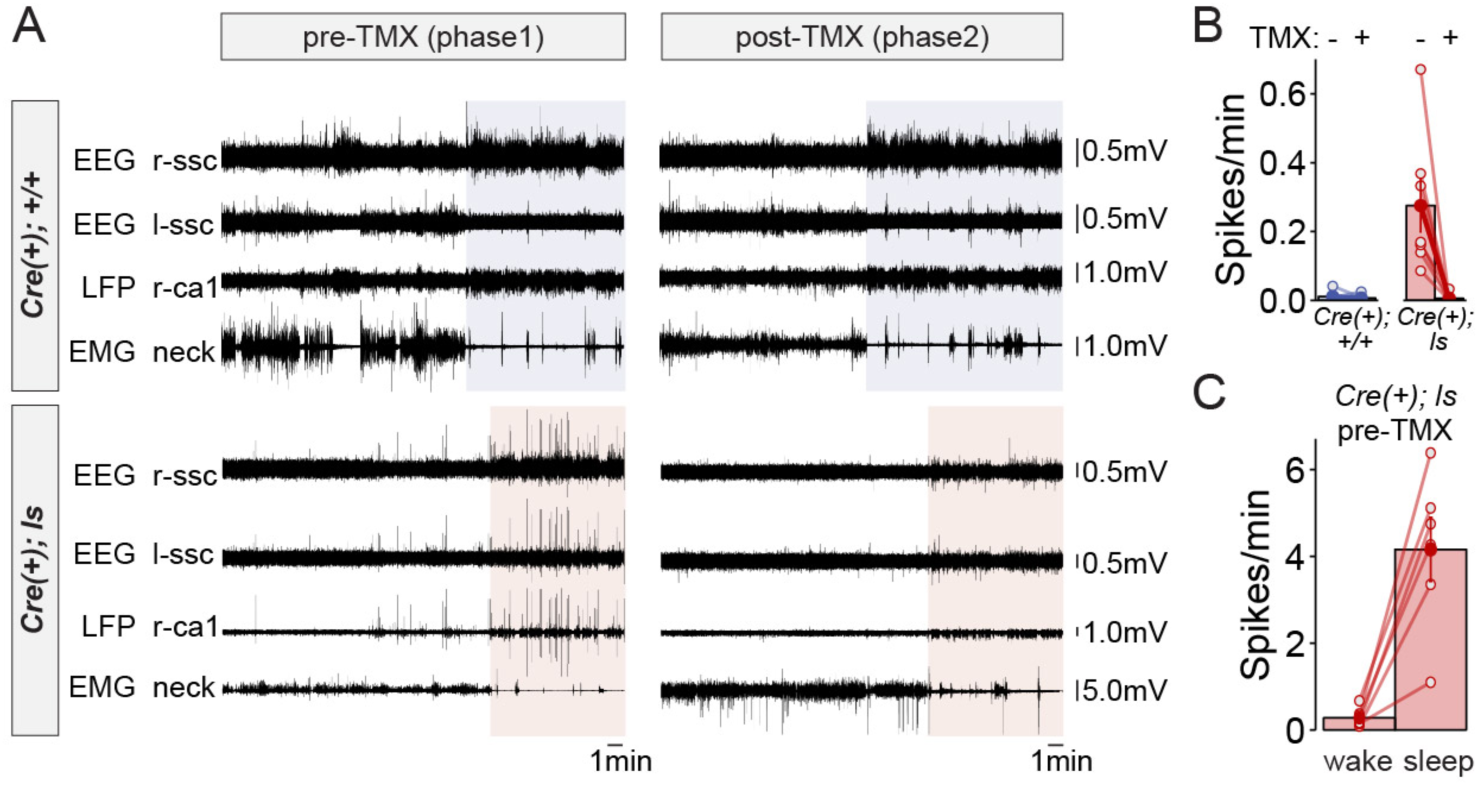
Rectification of state-dependent paroxysmal spiking events in *Syngap1* mutants after adult-initiated gene therapy. (A) Representative EEG/LFP traces from a WT [Cre(+); +/+] and *Syngap1* heterozygous mutant mouse [Cre(+); +/ls]. After initial recordings (pre-TMX), all animals were injected with tamoxifen. Post-TMX recordings were acquired 30 days after the last TMX injection. TMX rescued low levels of SynGAP protein in +/*Is* animals (see supplemental Figure 1). Highlighted areas correspond to periods of sleep (see methods). Phase I and Phase II recordings are from the same animals. **(B)** Frequency of spiking events observed in the hippocampal LFP channel during the wake phase (i.e. non-highlighted areas in panel A) from both pre- and post-tamoxifen recording sessions in each animal. Two-way repeated measure ANOVA. Main genotype effects: F(1,11)=10.1, p=.00879, Main TMX effects: F(1,11)=12.088, p=0.00518. Interaction between genotype and TMX: F(1,11)=9.777, p=.00963. Cre(+);+/+ n= 6, Cre(+);ls n= 7. **(C)** Comparison of the spiking frequency from the *h*LFP channel in Cre(+);ls mice during wake and sleep before Tamoxifen injections, paired-*t* test t(5)=-5.6007, p=0.002507 (n=5).

EEG waveforms are emerging as potential endpoints in clinical research studies for quantifying the efficacy of novel treatments for brain disorders [32, 33]. We were therefore interested in determining if EEG-recorded IIS events in *Syngap1* mice had parallels in humans expressing pathogenic *SYNGAP1* variants. While IIS events are commonly observed in both humans and animal models expressing genetic risk variants linked to epilepsy, worsening of these events during sleep is uncommon and is a symptom of a distinct cluster of epilepsy syndromes, such as electrical status epilepticus in sleep (ESES)/continuous spikes and waves during slow sleep (CSWS) [34]. Therefore, because sleep-dependent worsening of IIS events was observed in *Syngap1* mice, we searched for evidence of sleep-dependent worsening of paroxysmal spiking events in MRD5 patients. To do this, we mined data from a *SYNGAP1* patient registry, which has been used to uncover previously unreported phenotypes in this NDD population[35]. In this database, we found 12 entries that included a relatively complete medical history, including an

EEG report from a neurologist and a detailed genetic report defining the *SYNGAP1* variant. Each of the selected patients were confirmed to have a severe *SYNGAP1* variant expected to cause genetic haploinsufficiency (i.e. heterozygous knockout caused by either a nonsense variant or a frameshift caused by an InDel). In each of these unique patient entries, there were clear references to IIS activity **(Table 1)**. However, there were six unique entries where the neurologist noted that spike-related events worsened during sleep. In one patient (#5569), the neurologist noted that epileptiform activity was nearly constant during light sleep. In another (#1029), IIS was observed in 30% of recorded sleep. These clinical observations indicate that some patients with *SYNGAP1* haploinsufficiency have worsening EEG profiles during sleep, which strengthens the validity of sleep-dependent worsening of IIS in *Syngap1* mouse models.

Hippocampal oscillations recorded from LFP electrodes are linked to a range of cognitive functions, including various mnemonic process [36]. Because SynGAP re-expression in adult mutants improved remote contextual memory, we were interested in determining how this treatment impacted hippocampal oscillations, such as theta rhythms, that are thought to promote mnemonic processes. To do this, we investigated how SynGAP re-expression affected the amplitude of memory-linked oscillations recorded from the hippocampal LFP electrode before and after TMX injections. Theta rhythms were clearly observable in WT mice during both recording phases **(Fig. 4A)**. However, in recordings from Lox-Stop mice animals during Phase I, theta rhythms often appeared less robust during periods of free exploration compared to WT littermates **(Fig. 4B)**. After SynGAP re-expression, theta oscillations in these animals increased compared to pre-injection baseline recordings **(Fig. 4B)**. Across the entire Lox-Stop population, we found that theta-range signal amplitude was significantly enhanced by raising SynGAP protein levels in this population of adult Lox-Stop mice **(Fig. 4C)**, which is a finding consistent with improved performance in memory tasks considering that increased theta has been shown to correlate with better memory performance.

**Figure 4:**
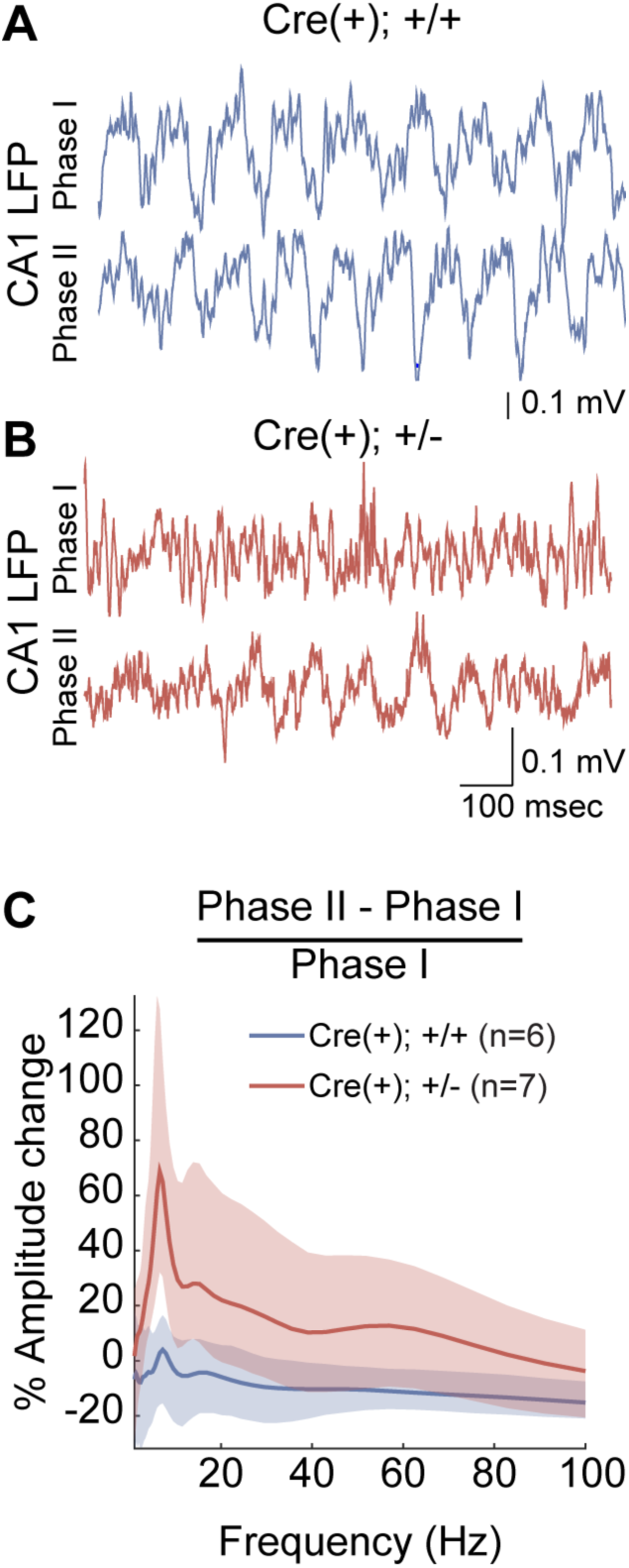
Increased amplitude of theta oscillations after SynGAP re-expression in adult *Syngap1* mutant mice. (A-B) CA1 LFP traces from a WT (A) and a *Syngap1* mutant (B) mouse during Phase I and Phase II sessions. C) Grand average of within-subjects changes in signal amplitude across the full spectrum of hippocampal rhythms. The amplitude change was normalized by the average amplitude during Phase I sessions. The shaded areas represent 95% bootstrapped confidence intervals. Significant increases in amplitude in Phase II were detected in the 6-12 Hz theta range (Permutation test: p = 0.0128, 5000 shuffles).

Theta rhythms in mice are modulated by running speed [37] and it has been reported that *Syngap1* Het mice are hyperactive in some, but not all, contexts [23]. Therefore, we tracked locomotor activity in all recorded mice across both sessions to determine if phase-specific changes in activity levels may have contributed to changes in Theta amplitude **(Supplemental Figure 3)**. Similar to past studies, Cre(+) Lox-Stop Het mice were hyperactive compared Cre(+)WT controls because there was a main effect of genotype on distance traveled. However, we did not observe an effect of phase or an interaction between phase and genotype in this analysis. Therefore, the phase-independent increases in locomotor activity in Het mice is unlikely to explain the phase-specific changes to theta rhythms in these animals.

## Discussion

The principal finding in this study is that genetic reversal of *Syngap1* pathogenicity in adult mice improves deficits in memory, excitability and hippocampal/cortical function. On the surface, these exciting results are somewhat paradoxical given that we have previously found that *Syngap1* pathogenicity causes hardwired impairments in the development of forebrain circuits that mediate cognitive functions [22, 24, 38]. These past studies led to the conclusion that impaired assembly of forebrain circuits contributed to life-long impairments in brain function because a set of behaviors were resistant to adult re-activation of SynGAP protein expression. How then can these current findings be reconciled with past findings demonstrating that certain behavioral paradigms are not sensitive to adult re-expression of SynGAP protein? The most parsimonious explanation is that not all behavioral impairments common to the *Syngap1* endophenotype are governed by hardwired circuit changes, which in turn reflect altered neurodevelopmental processes. Some behavioral impairments may be caused by altered "real-time" neuronal signaling that is not restricted to defined periods of neurodevelopment[5]. Elegant work in Shank3 mouse models directly supports this interpretation[39]. Feng and colleagues demonstrated that adult reversal of *Shank3* pathogenicity improved some behavioral domains, while others remained completely impaired even though protein levels were restored to WT levels. Restoring Shank3 protein levels early in development prevented the emergence of adult reversal-resistant behavioral impairments, indicating that low expression of germline Shank3 protein disrupts both developmental and non-developmental cellular processes.

There is evidence in the literature that *Syngap1* may have unique functions during both development and adulthood, which supports an evolving interpretation of SynGAP function in the brain. For instance, *Syngap1* heterozygosity causes a transient increase in baseline postsynaptic function in CA1 Schaffer collateral synapses during postnatal development[22]. This is a true developmental impact of *Syngap1* pathogenicity because inducing pathogenicity in adulthood does not induce a change in baseline excitatory synapse function in hippocampal synapses. Interestingly, baseline excitatory function of this pathway normalizes by adolescence, as evidenced by several groups[40] [41], including our own[22], reporting normal postsynaptic baseline function in this synapse in mature animals. However, there is a dramatic impairment in LTP stability at these synapses in adult *Syngap1* mice[40, 41]. This function of *Syngap1* appears unrelated to neurodevelopment because LTP can be completely rescued after adult re-expression of low SynGAP protein levels[26]. Thus, *Syngap1* exhibits both developmental and non-developmental functions with the same synapses in the hippocampus. As a consequence, not all deficits related to neuronal function in the adult *Syngap1* Het mouse brain can be attributed to hardwired circuit damage caused by impaired neurodevelopment.

The real-world impact of our current findings is two-fold. First, therapies that improve the expression and/or function of SynGAP protein may be beneficial to MRD5 patients of all ages. Early therapeutic intervention should remain the primary NDD treatment strategy because this would in theory protect the brain from hardwired circuit damage caused by disruptions to critical neurodevelopmental processes. However, adult re-expression of SynGAP appears to correct neurophysiological imbalances within circuits that predispose the brain to seizure generation and certain forms of cognitive impairment. Therefore, the impact of SynGAP-raising therapies on adult patients could be significant. 1t remains unclear if adult re-expression of SynGAP corrects seizure susceptibility and memory impairments through a common mechanism. Future studies will be necessary to determine the molecular and cellular mechanisms that are recovered in the forebrain after SynGAP re-expression. Moreover, it will be critical to link these recovered molecular and cellular processes to circuit-level processes that directly contribute to behavioral memory and seizure. This type of experimental strategy may help to elucidate novel neural circuit correlates of behavioral alterations and excitability impairments associated with NDDs.

The second real-world implication of this study is that it may have identified a candidate translatable biomarker that can serve as an endpoint for efficacy testing of novel treatments for MRD5 patients. Our data indicate that IIS events appear to worsen during sleep in both humans and mice with *SYNGAP1/Syngap1* haploinsufficiency. Sleep-linked worsening of IIS is a hallmark of epilepsy syndromes with significant cognitive impairment [34]. We found that SynGAP re-expression in adult mutant mice eliminated these pathological events, which are also detected by EEG electrodes, during both wakefulness and sleep. Considering that this treatment strategy also improved measures of behavioral seizure and memory, the severity of IIS during sleep may be a predictor of generalized cortical impairment associated with *SYNGAP1* pathogenicity. If a relationship between the severity of state-dependent IIS and cognitive impairment was observed in patients, then these signals may be useful as biomarkers for improved cortical function in translational studies aimed at identifying effective therapeutic strategies for *SYNGAP1* patients.

## Methods

### Animals

*Syngap1* constitutive *(Syngap1*^+/-^) and reversal *(Syngap1*^+/ls^) mice were constructed and maintained on a mixed genetic background (C57BL/6j:129/SvEv) as previously described^17^. Inducible CAGG-Cre-ER^TM^ male mice were purchased from Jackson Laboratories (JAX stock #004682) and crossed to female *Syngap1*^+/ls^ for adult reversal studies. Male and female experimental mice were utilized for all studies. Animals were group housed (n=4/cage) by sex and otherwise randomly assigned to cages yielding mixed genotypes. Animals were kept on a normal light-dark cycle and had free access to food and water. Animal husbandry and experiments were conducted according to TSRI Institutional Animal Care and Use Committee code.

### Cre Induction

TMX (Sigma T5648, St. Louis, MO) was prepared in corn oil containing 10% EtOH to a final TMX dosage of 100 mg/kg, injectable concentration of 20 mg/ml, and volume of 5 ml/kg and administered (intraperitoneal) once a day for five consecutive days starting at PND60.

### Behavior

Mice (PND90-120) were handled for several minutes on three separate days prior to commencement of behavioral testing. Tails were marked for easy identification and access from home cages during testing. Experimenters were blind to genotype while conducting all tests. Various cohorts of mice were utilized consisting of one cohort of constitutive *Syngap1*^*+/+*^ and *Syngap1*^*+/-*^ mice and one cohort of *Syngap1*^*+/Is*^ mice in obtaining results for Figure 1 as well as two combined cohorts each for TMX(-) and for TMX(+) inducible Cre *Syngap1*^*+/+*^ and *Syngap1*^*+/Is*^ mice for generating results for Figure 1E&F and Figure 2. Separate cohorts were run for TMX-inducible Cre *Syngap1*^*+/+*^ and *Syngap1*^*+/Is*^ mice for the flurothyl-induced seizure test and Western blot analysis. Several mice from the TMX(+) inducible Cre cohorts died prior to ASR/PPI and seizure testing. Samples sizes for *Syngap1*-related studies in our lab have been estimated using GPower 3.0. Estimates for effect size and variance of behavioral data were based on published studies using the *Syngap1* mouse line. Experimenters were blind to genotype while conducting behavioral analyses.

### Contextual fear conditioning

A dedicated fear conditioning room in the mouse behavior core contained four fear conditioning apparati that can be used in parallel. Each apparatus is an acrylic chamber measuring approximately 30 x 30 cm (modified Phenotyper chambers, Noldus, Leesburg, VA). The top of the chamber is covered with a unit that includes a camera and infrared lighting arrays (Noldus, Ethovision XT 11.5, Leesburg, VA) for monitoring of the mice. The bottom of the chamber is a grid floor that receives an electric shock from a shock scrambler that is calibrated to 0.40 mA prior to experiments. The front of the chamber has a sliding door that allows for easy access. The chamber is enclosed in a sound-attenuating cubicle (Med Associates) equipped with a small fan for ventilation. Black circular, rectangular and white/black diagonal patterned cues were placed outside each chamber on the inside walls of the cubicles for contextual enhancement. A strip light attached to the ceilings of the cubicles provided illumination. A white noise generator (~65 dB) was turned on and faced toward the corner of the room between the cubicles. The fear conditioning paradigm consisted of two phases, training, followed by testing 1 and 26, or 30d thereafter. The 4.5 min training phase consisted of 2.5 min of uninterrupted exploration. Two shocks (0.40 mA, 2 s) were delivered, one at 2 min 28 s, the other at 3 min and 28 s from the beginning of the trial. During testing, mice were placed into their designated chambers and allowed to roam freely for 5 min. Immobility durations (s) and activity (distances moved (cm)) during training and testing were obtained automatically from videos generated by Ethovision software. Activity suppression ratio levels were calculated: 0-2 min activity during testing/(0-2 min activity during training + testing).

### Flurothyl-induced seizures

Flurothyl-induced seizure studies were performed based on prior studies with some modifications[22, 26, 42]. Briefly, experiments were conducted in a chemical fume hood. Mice were brought to the experimental area at least 1 h before testing. To elicit seizures, individual mice were placed in a closed 2.4-L Plexiglas chamber and exposed to 99% Bis (2,2,2-triflurothyl) ether (Catalog# 287571, Sigma-Aldrich, St. Louis, MO,). The flurothyl compound was infused onto a filter paper pad, suspended at the top of the Plexiglas chamber through a 16G hypodermic needle and tube connected to a 1ml BD glass syringe fixed to an infusion pump (KD Scientific, Holliston, MA, USA, Model: 780101) at a rate of 0.25 ml/min. The infusion was terminated after the onset of a hind limb extension that usually resulted in death. Cervical dislocation was performed subsequently to ensure death of the animal. Seizure thresholds were measured as latency (s) from the beginning of the flurothyl infusion to the beginning of the first myoclonic jerk (1st clonic), then to generalized tonic/clonic seizure (T/C), and finally to total hind limb extension (THE). Experimenters were blind to genotype while conducting seizure threshold analyses.

### lmmunoblotting

Western blot analysis was performed on protein lysates extracted from the hippocampi of adult mice and dissected in ice-cold PBS containing Phosphatase Inhibitor Cocktails 2 and 3 (Sigma-Aldrich, St. Louis, MO) and Mini-Complete Protease Inhibitor Cocktail (Roche Diagnostics) and immediately homogenized in RIPA buffer (Cell Signaling Technology, Danvers, MA), transferred to tubes in dry ice, and stored at −80°C. Sample protein concentrations were measured (Pierce BCA Protein Assay Kit, Thermo Scientific, Rockford, IL), and volumes were adjusted to normalize microgram per microliter protein content. 10µg of protein per sample were loaded and separated by SDS-PAGE on 4-15% gradient stain-free tris-glycine gels (Mini Protean TGX, BioRad, Hercules, CA), transferred to low fluorescence PVDF membranes (45µ) with the Trans-Blot Turbo System (BioRad). Membranes were blocked with 5% powdered milk in buffer and probed with pan-SynGAP (1:10,000, PA1-046, Pierce/Thermo Scientific) overnight at 4°C and HRP-conjugated anti-rabbit antibody (1:2,500, W4011, Promega) for 1hr at room temperature followed by ECL signal amplification and chemiluminescence detection (SuperSignal West Pico Chemiluminescent Substrate; Thermo Scientific, Rockford, IL). Blot band densities were obtained using the Alpha View imaging system (Alpha Innotech). SynGAP levels of immunoreactivity were assessed by densitometric analysis of generated images with ImageJ. Values were normalized to total protein levels obtained from blots prior to antibody incubations.

### Video-EEG recordings

All research and animal care procedures were approved by the Baylor College of Medicine Institutional Animal Care and Use Committee. Experiments were carried out on 8 Cre(+)/Syngap1 Lx-ST heterozygous mice and 8 littermate Cre(+)/Syngap1 wild type controls. Experimenters were blind to the genotypes. Animals were bred at The Scripps Research Institute and experimental mice transferred to Baylor College of Medicine at ~4-5 weeks of age. Animals at 11 weeks of age were secured on a stereotaxic frame (David Kopf) under 1-2% isoflurane anesthesia. Each mouse was prepared under aseptic condition for the following recordings: Teflon-coated silver wires (bare diameter 127 µm, A-M systems) were implanted bilaterally in the subdural space of the somatosensory cortex[43] (0.8 mm posterior, 1.8 mm lateral to the bregma, reference at the midline over the cerebellum) for cortical EEG as well as in the neck muscles for electromyogram recordings to monitor mouse activity. An additional electrode constructed with Teflon-coated tungsten wire (bare diameter 50 µm, A-M systems) was sterotaxically implanted in the CA1 of the hippocampus[43] (1.9 mm posterior, 1.0 mm lateral, and 1.3 mm below the bregma) with reference in the ipsilateral corpus callosum for local field potential recordings. All of the electrode wires together with the attached miniature connector sockets were fixed on the skull by dental cement. After 2 weeks of post-surgical recovery, mice received Phase I video-EEG recordings (2 hours per day for 3 days). Signals were amplified (100x) and filtered (bandpass, 0.1 Hz - 1 kHz) by 1700 Differential AC Amplifier (A-M Systems), then digitized at 2 kHz and stored on disk for off-line analysis (DigiData 1440A and pClamp10, Molecular Devices). The time-locked mouse behavior was recorded by ANY-maze tracking system (Stoelting Co.). In addition, manual ON/OFF camcorder was used to monitor the behavior at higher resolution. From the day after Phase I recordings of video-EEG, all of the mice received daily injections of Tamoxifen *(as above)* for 5 days. One month later, animals went through Phase II video-EEG recordings (three 2-hour sessions over 3 days) under the same settings as in Phase I. At the end of the experiments, mice were euthanized, and the hippocampi were dissected to determine the efficacy of SynGAP protein re-expression.

### High-Amplitude lnterictal Spike Quantification

Axograph3 or pClamp10 software (Molecular Devices, San Jose, CA) was used to detect high-amplitude spiking events. The thresholded was set at +1mV in the CA1 depth channel for all animals and all events that exceeded the threshold were logged. Events were occasionally rejected as "non-physiological". These rejected signals were identified by their usually high signal amplitude and time-locked peaks in more than one channel and their atypical shape compared to paroxysmal spikes. Behavioral epochs were segregated into sleep and wake phases. Sleep was inferred from an abrupt quieting of the EMG signal and confirmation of immobility from time-locked videos. Experimenters were blind to genotype while conducting spike quantification analyses.

### Analysis of hippocampal oscillations

Analyses focused on the first 10 minutes of each recording session for each mouse to account for experience-dependent changes in mouse hippocampal rhythms [37]. Videos of mouse behavior and EMG recordings were used to verify that mice were engaged in active locomotion during this period, considering that theta oscillations occur most prominently in the hippocampus during voluntary movement [44]. Time-resolved amplitudes of CA1 LFP recordings were estimated using a complex Morlet wavelet transform with a width parameter of six periods, evaluated at 50 frequencies logarithmically spaced between 1 and 100 Hz [45]. Wavelets were normalized such that their area under the curve was equal to one [46]. The magnitude of theta (6- 12 Hz) oscillations across time was quantified using the ratio of theta to delta (1-4 Hz) power. The power ratio was smoothed by a moving average time window of 1 second. The one-second time window with the highest theta-delta power ratio for each mouse in each Phase was selected for the example CA1 LFP recordings shown in Figure 4A & B and Supplementary Figure 4A & B. To account for the variability of theta amplitudes across mice (see Supplementary Figure 4A & B), amplitude changes between Phase I and Phase II sessions were calculated within each mouse and then normalized by the corresponding amplitude during Phase I sessions (Figure 4C). Experimenters were blind to genotype while conducting LFP analyses.

### Collection and analysis of data from the Retrospective SYNGAP1 Natural History Study Registry

The *SYNGAP1* Patient Registry (https://syngap1registry.iamrare.org) is funded through the National Organization of Rare Disorders. This study was approved through the Hummingbird Institutional Review Board and meets all relevant ethical regulations for protections for human subjects. It is actively managed by a board of trustees comprised of a team of seven stakeholders, including parents with affected children, clinician-scientists that care for MRD5 patients, and neurobiologists that study the gene. The *SYNGAP1* (MRD5) Natural History Study Registry is a retrospective longitudinal web-based observational natural history study. Parents or guardians provided informed consent prior to depositing medical history data into the registry. Participants with *SYNGAP1* (MRD5) will be followed throughout the course of their lives with either the participant or authorized respondents contributing data at varying intervals throughout the course of the study. Initially, when a new patient is registered, data is collected on demographics, quality of life, medical history including genetic reports, disease phenotypes, event episodic data, retrospective data, participant review of systems, medication and diagnostic data. Each registrant is given a unique identifier to facilitate anonymization of patient data. Initial data collection is done through a series of questionnaires, including a survey of sensory and sensory-related issues. The structure of the database and all questionnaires were reviewed and approved by the members of the Board of Trustees.

To acquire information about EEG during sleep and wakefulness in the *SYNGAP1* patient population, the registry database was queried for all entries that: 1) had a detailed genetic report that confirmed the presence of a severe *SYNGAP1* loss-of-function variant likely to induce genetic haploinsufficiency; 2) had at least a narrative report from a neurologist that discussed the findings of an EEG study. **Table 1** summarizes each patient-specific *SYNGAP1* variant and findings from their EEG study. A subset of EEG reports contained information about the patient's overall clinical presentation and medications at the time of the EEG study. This information was included in Table 1 where appropriate.

### Statistics

Statistical tests were conducted using SPSS and R packages importing Excel files containing raw data. Based on our prior experience with these behaviors a general parametric statistical approach was used to assess data sets. Unpaired t tests for experiments with two groups comparing genotypes and two-factor ANOVAs for experiments with four groups assessing contingencies of Cre and genotype factors were performed to assess activity suppression ratio values associated with fear conditioning, ASR and habituation to ASR, and immunoblot data. MANOVAs were performed to assess Cre/genotype contingencies for the multiple levels of factors associated with seizure threshold (1^st^ clonic, T/C, and THE) and PPI (PPI4, PPI8, and PPI16). Explorations of data sets showed reasonably normal distributions and homogeneity of variances as assessed by Levene's test for equality of variances as well as Box's test for equality of covariance matrices except with that in Fig1E and in Fig1B where we employed the Wilcoxon rank sum test. In general, we did not systematically search for outliers in our data, yet we did observe one instance of an anomalous value occurring in figure 1C (animal #881) that had an ASR value of >0.5. Values >0.5 in this test are highly unusual and this data point was far removed from the others. A Grubbs' Test for Outliers flagged the value. It was therefore removed from the experiment. A nonparametric permutation test (5000 shuffles) was used to assess differences in oscillatory activity (Figure 4).

---

**Supplemental Figures**

**Supplemental Figure 1:**
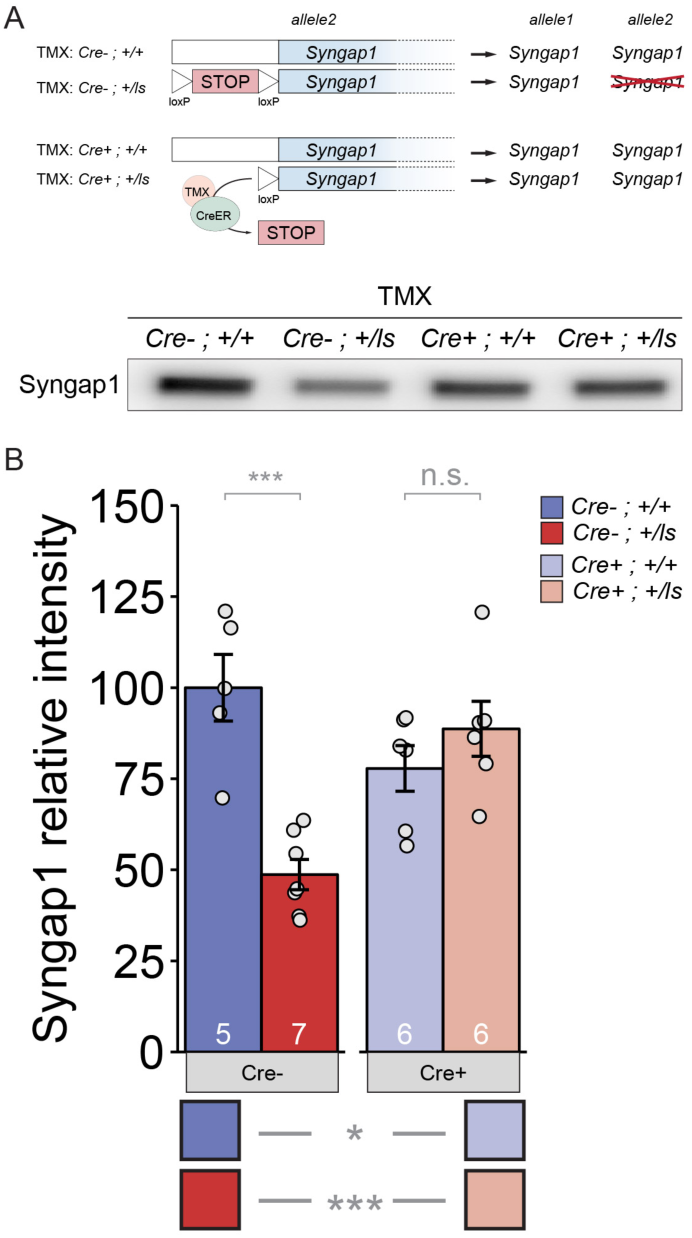
TMX-induced restoration of SynGAP protein levels in adult *Syngap1*^*Cre+;+/Is*^ mice. (A) Western blot demonstrating expression levels of total SynGAP in Cre(-) or Cre(+) heterozygous Lox-Stop mice and WT littermates. **(B)** Densitometric analysis of SynGAP. band intensities were normalized to total protein levels and transformed to % of the *Syngap1*^*Cre+;-/Is*^ group mean. Two-factor ANOVA. Main effects: Cre p=.198, Genotype p=.007, Interaction p= 1.554E-4. Pairwise comparisons: *Syngap1*^*Cre-;+/+s*^ vs *Syngap1*^*Cre-;+/Is*^ p=2.853E-5; *Syngap1*^*Cre+;+/+*^ vs *Syngap1*^*Cre+;+/Is*^ p=.261; *Syngap1*^*Cre-;+/+*^ vs *Syngap1*^*Cre+;+/+*^ p=.036; *Syngap1*^Cre-;+;+/Is^ vs *Syngap1* p=2.642E-4.

**Supplemental Figure 2:**
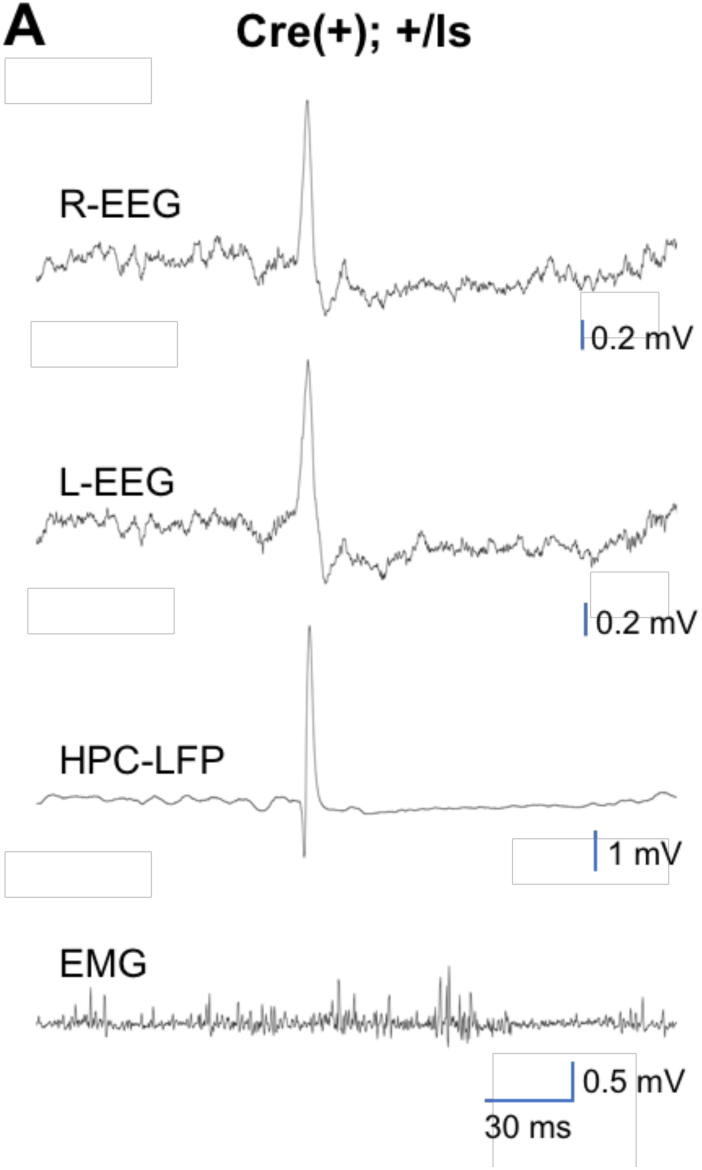
Generalization of high-amplitude spikes across the forebrain. **(A)** Representative traces from all channels during a Phase I recording from a Cre(+) Lox-Stop mouse.

**Supplemental Figure 3:**
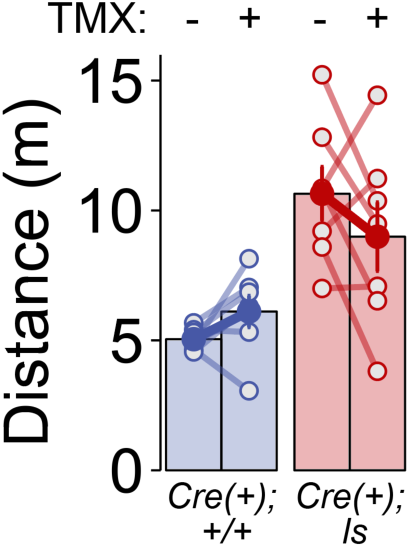
Effect of genotype, but not phase, on horizontal activity during neurophysiological recordings. Cre(+) WT and Cre(+) Lox-Stop mice were video tracked for distances traveled during the first ten minutes of recording during Phase I (TMX-) and Phase II (TMX+) sessions. RMANOVA-Group: F(1,12)=16.527, p=.002; Phase: F(1,12)=.164, p=.692; Group x Phase: F(1,12)=3.521, p=.085.

**Supplemental Figure 4:**
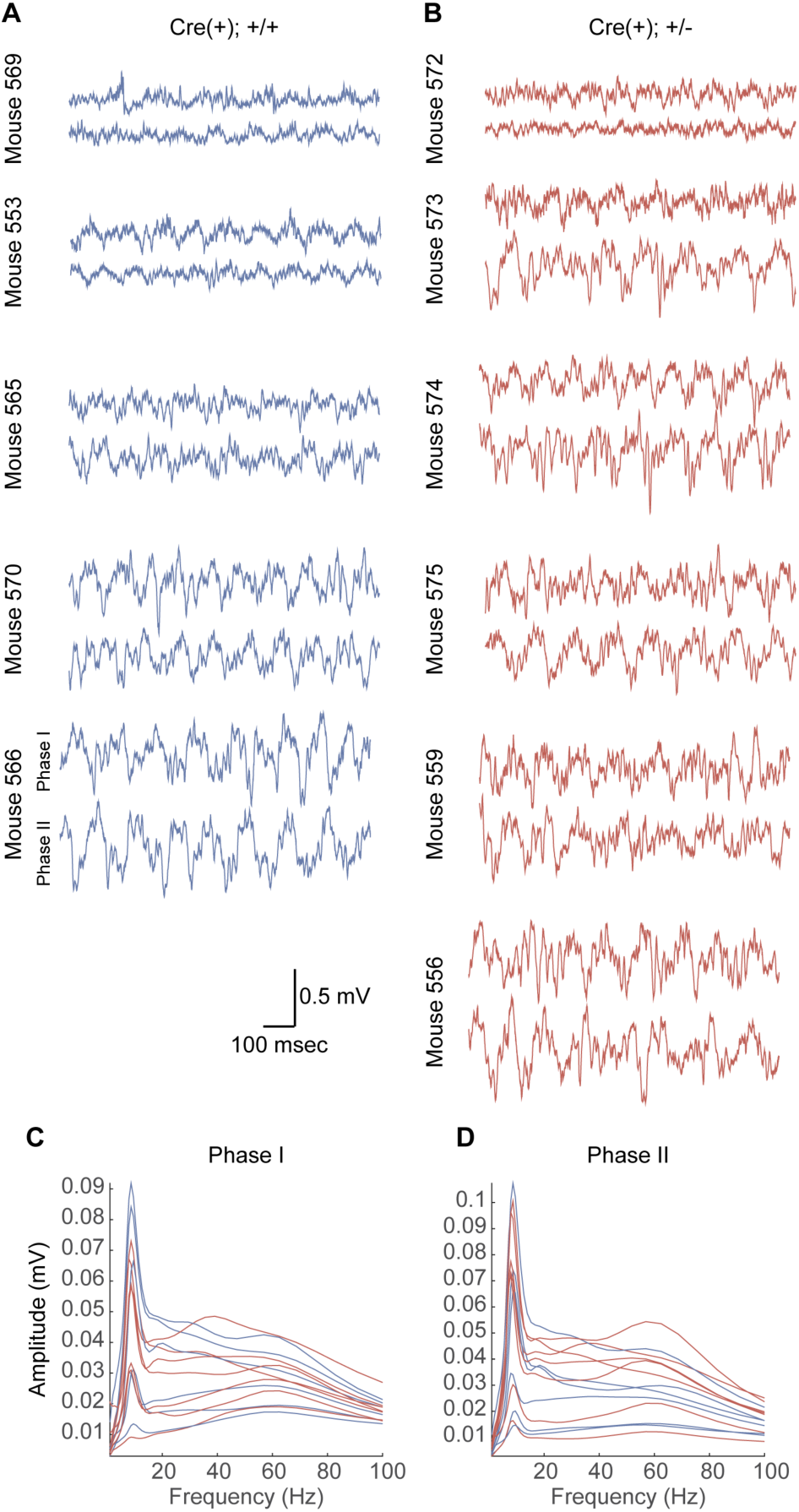
Amplitude of theta oscillations in each mouse during Phase I and Phase II recording sessions. (A-B) CA1 LFP recordings from WT (A) and *Syngap1* mutant (B) mice during Phase I and Phase II sessions. (C-D) Average amplitude spectra for each mouse during Phase I (C) and Phase II (D) sessions. Individual mice are indicated with individual lines, and WT and *Syngap1* mutant spectra are depicted in blue and red, respectively.

**Author Contributions**
TKC, CR, JT and MK performed experiments. TKC, CR, JT, LLC, CAM, and GR designed experiments. TKC, CR, MK, TV, JLH, EH, CAM, and GR analyzed data. GR wrote the manuscript and conceived the study. TKC, CR JLH, LLC, EH and CAM edited the manuscript.

## Acknowledgements

This work was supported in part by NIH grants from the National Institute of Mental Health [MH096847 and MH108408 (GR), MH105400 (CAM), MH102450 (LLC)], the National Institute for Neurological Disorders and Stroke [NS064079 (GR)], and the National Institute for Drug Abuse [DA034116 and DA036376 (CAM), T32DA01892 (EH)]. JLH is supported by a National Institute for Neurological Disorders and Stroke Mentored Clinical Scientist Research Career Development Award [NS091381] and the Robbins Foundation.

The video-EEG experiments were performed by the IDDRC Neuroconnectivity Core at Baylor College of Medicine.

